# Distal Gene Expression Governed by Lamins and Nesprins via Chromatin Conformation Change

**DOI:** 10.1101/2025.04.01.646570

**Authors:** Haihui Zhang, Zhengyang Lei, Fatemeh Momen-Heravi, Peiwu Qin

**Affiliations:** Institute of Biopharmaceutical and Health Engineering, Shenzhen International Graduate School, Tsinghua University, Shenzhen, Guangdong, China; Department of Orofacial Sciences, University of California San Francisco, San Francisco, CA, USA

**Keywords:** RNA-seq, Lamin A, *LMNA*, Nesprin, *SYNE2*, dCas9, telomere live-cell imaging

## Abstract

The nuclear lamina is a vital structural component of eukaryotic cells, playing a pivotal role in both physiological processes, such as cell differentiation, and pathological conditions, including laminopathies and cancer metastasis. Lamina associated proteins, particularly lamins and nesprins, are integral to mechanosensing, chromatin organization, and gene regulation. However, their precise contributions to gene regulation remain incompletely understood. This study explores the functions of lamin A, *LMNA*, and *SYNE2* in gene expression, with a particular focus on their influence on distal chromatin interactions and conformational changes. Using inducible shRNA knockdown, RNA-seq analysis, and dCas9-mediated live imaging of chromosomes, we demonstrate that lamin A affects RNA synthesis, *LMNA* governs chromatin spatial organization, and *SYNE2* regulates chromatin modifications. Furthermore, both lamins and nesprins enhance telomere dynamics. These findings elucidate nuclear envelope-associated mechanisms in gene regulation, offering valuable insights into chromatin dynamics under both physiological and pathological contexts.

## 1. Introduction

The three-dimensional organization of the genome within the nucleus plays a crucial role in coordinating gene expression, ensuring the stable repression of heterochromatin while permitting the dynamic regulation of transcriptionally active regions. This organization is intimately connected to the nuclear envelope, a specialized membrane enriched with proteins that actively contribute to chromatin architecture and gene regulation^1^. Among these proteins, the nuclear lamina-a filamentous meshwork of intermediate filament (IF) proteins known as lamins—forms a structural scaffold that anchors chromatin to the nuclear periphery, thereby influencing nuclear mechanics and genomic stability ^2^. Lamins interact with the linker of nucleoskeleton and cytoskeleton (LINC) complex to form a direct mechanotransduction bridge that links the cytoskeleton to the extracellular matrix (ECM) while simultaneously influencing chromatin topology. A-type lamins, encoded by the *LMNA* gene, include lamin A and lamin C, whereas B-type lamins (*LMNB1* and *LMNB2*) maintain constitutive roles in nuclear structure across different cell types ^3–7^.

Lamins have been implicated in diverse biological processes, including stem cell differentiation, cancer progression, aging, and laminopathies, underscoring their multifaceted role in nuclear integrity and genome regulation. Nesprins, giant spectrin-repeat proteins of the LINC complex, serve as critical linkers between the cytoskeletal filaments and the nuclear lamina, providing structural integrity and modulating intracellular signaling ^8,9^.

While significant progress has been made in understanding the role of lamins in genome organization, the precise mechanisms by which lamins and nesprins regulate gene expression through distal chromatin interactions remain incompletely understood ^10,11^. Notably, recent evidence suggests a reciprocal interplay between transcription and chromatin conformation, where gene activity can influence chromatin folding and vice versa ^12^. However, whether lamins and nesprins actively govern chromatin remodeling and isoform switching beyond their well-characterized functions in mechanotransduction remains an open question.

Lamina-associated domains (LADs) represent large chromatin regions tethered to the nuclear lamina and are generally transcriptionally repressive, enriched with heterochromatic marks such as H3K9me2/3 and H3K27me3. These domains play a pivotal role in nuclear architecture by segregating active and inactive chromatin compartments. Disruptions in LAD organization have been implicated in diseases such as cancer and laminopathies, highlighting the importance of nuclear envelope components in maintaining genomic stability. However, the extent to which lamins and nesprins regulate isoform switching and chromatin dynamics remain largely unexplored.^13,14^

This study aims to dissect the roles of lamin A, *LMNA*, and *SYNE2* in gene expression and chromatin organization, focusing on their influence on distal chromatin interactions and isoform switching. Using inducible shRNA-mediated knockdown, RNA sequencing (RNA-seq), and dCas9-based live-cell imaging of chromosomes, we systematically investigate their contributions to chromatin conformation and transcriptional regulation. Our findings suggest that lamin A contributes to RNA synthesis, supports chromatin spatial organization through *LMNA*, and that *SYNE2* influences chromatin modifications as reflected in transcript levels. Furthermore, we demonstrate that both lamins and nesprins impact telomere dynamics, linking nuclear envelope function to chromatin behavior.

By uncovering distinct, isoform-specific and spatially segregated transcriptional responses, our study provides new mechanistic insights into how nuclear lamina components regulate distal gene expression. These findings have broad implications for understanding nuclear architecture in both physiological and pathological contexts, including cancer, aging, and laminopathies.

## 2. Materials and methods

### 2.1 Cell culture

Human osteosarcoma (U2OS) cells (#HTB-96, ATCC, USA) and human embryonic kidney (HEK293T) cells (#CRL-3216, ATCC, USA) were cultured in modified Dulbecco’s Modified Eagle’s Medium (#11965092, DMEM; Gibco), supplemented with 10% fetal bovine serum (FBS). U2OS cells were cultured using HyClone™ characterized tetracycline-screened FBS (#SH30071.03T) at a density of 5 × 10^4^ cells per cm^2^ for RNA extraction experiments, and at a density of 1 × 10^3^ cells per cm^2^ for Chromosomes live imaging experiments. While HEK293T cells were cultured with Gibco FBS (#A5256701) at an initial density of 5 × 10^4^ cells per cm^2^. The medium also contained 100 U ml^-1^ penicillin and 100 µg ml^-1^ streptomycin (#15140122, Gibco). All cultures were maintained in a humidified incubator at 37 °C with 5% CO₂.

### 2.2 Construction of stable U2OS cells within inducible shRNA plasmids

To generate doxycycline-inducible stable knockdown cell lines targeting scramble control, lamin A, *LMNA*, and *SYNE2*, specific short hairpin RNAs (shRNAs) were employed. Lamin A and lamin C are two isoforms produced by alternative splicing of the *LMNA* gene. While they share an identical N-terminal sequence, they differ in their C-termini, resulting in distinct biochemical properties and functional roles in nuclear structure and gene regulation. The shRNA against lamin A (shLaminA) targets the 3`′` untranslated region (UTR) of the *LMNA* gene, specific to prelamin A, which is post-translationally processed into mature lamin A. The shRNA designated as shLMNA targets a region within the coding sequence of *LMNA* that is shared by both lamin A and lamin C, corresponding to amino acids 122–129 (KKEGDLIA) of lamin A/C (RefSeq: NM_001406985.1). The shRNA against *SYNE2* (shSYNE2) targets a sequence encoding amino acids 5133–5140 (KRYERTEF) of the SYNE2 protein (RefSeq: NM_182914.3). Targeting shRNAs (**Supplementary Table S1**) were cloned into the pLKO-Tet-On (#21915, Addgene) inducible vectors by AgeI and EcoRI restriction enzymes (New England Biolabs, USA). HEK293T cells were transfected with pLKO-Tet-On-shRNA, psPAX2 and pMD2.G with ratio of 4:3:1 using Lipofectamine™ 3000 (#L3000015, ThermoFisher Scientific, USA). Lentiviral supernatants were harvested after 48 h, filtered through a 0.45-µm filter, and stored at −80 °C until further use. The Multiplicity of Infection (MOI) was determined by limiting dilution of lentiviral particles. U2OS cells were infected with lentivirus at an MOI of 0.8 in the presence of 8 µg ml^-1^ polybrene (#H9268, Hexadimethrine bromide; MilliporeSigma, USA). Following infection, cells were selected for five days using 4 µg ml^-1^ puromycin (#P4512, MilliporeSigma, USA). Single-cell clones were subsequently isolated and expanded in the presence of 2 µg ml^-1^ puromycin to establish doxycycline-inducible shRNA-knockdown stable cell lines. Multiple clones per shRNA were screened for knockdown efficiency by reverse transcription quantitative real-time PCR (RT-qPCR). A single clone with robust and consistent knockdown was selected for downstream analyses, including RNA-seq. For doxycycline-inducible shRNA-knockdown stable cell lines, the cells were treated with 100 ng ml^-1^ doxycycline (#D5207, MilliporeSigma, USA) for 48 hours.

### 2.3 RT-qPCR

The knockdown efficiency of targeting genes in the stable U2OS cells containing inducible shRNA plasmids was assessed using RT-qPCR. Total RNA was extracted by Trizol (#T3934, MilliporeSigma, USA), 500 ng of total RNA was used as a template for reverse transcription into cDNA using the High-Capacity cDNA Reverse Transcription Kit (#4374966, ThermoFisher Scientific, USA) and real-time PCR was performed using PowerUp™ SYBR™ Green Master Mix (#A25778, ThermoFisher Scientific, USA). For the procedure of real-time PCR, cDNA templates, SYBR, and primers were mixed and loaded to Applied Biosystems™ 7500 Real-Time PCR Systems, started with an initial denaturation at 95 °C for 300 seconds to separate DNA strands. Following 40 cycles of denaturation (95 °C for 20 seconds), annealing (55 °C for 20 seconds), and extension (72 °C for 20 seconds). The relative transcript abundance was initially normalized to *GAPDH* and subsequently adjusted relative to the no-doxycycline-treated controls using the ΔΔCt method. Primer sequences are shown in **Supplementary Table S2**.

### 2.4 Library preparation for transcriptome sequencing (RNA-seq)

Total RNA, including mRNA and lncRNA, was used to prepare sequencing libraries. RNA was purified from total RNA using probes to remove rRNA by Ribo-Zero rRNA Removal Kit (#15066012, Illumina, USA). RNA integrity and quantification were assessed using the Bioanalyzer 2100 system (Agilent Technologies, USA). RNA was fragmented with divalent cations at high temperature in a First Strand Synthesis Reaction Buffer (5×). First strand cDNA was synthesized with random hexamer primers and M-MuLV Reverse Transcriptase (RNase H), followed by second strand cDNA Synthesis using DNA Polymerase I and RNase H. The concentration of cDNAs in the library was quantified to 1 ng μl^-1^ using a Qubit 2.0 fluorometer. Overhangs were converted to blunt ends using exonuclease and polymerase activities. After adding adenylated 3’ ends to the DNA fragments, NEBNext Adapters with hairpin loops were ligated for hybridization.

To select cDNA fragments 370–420 bp in length, libraries were purified with the AMPure XP system (Beckman Coulter). Size-selected, adaptor-ligated cDNA was treated with 3 µl USER Enzyme (New England Biolabs) at 37 °C for 15 minutes, followed by 95 °C for 5 minutes. PCR amplification was performed using Phusion High-Fidelity DNA Polymerase, Universal PCR primers, and Index (X) primers. The PCR products were purified using the AMPure XP system, and library quality was assessed using the Agilent Fragment Analyzer 5400 system.

Clustering of index-coded samples was carried out on a cBot Cluster Generation System using the TruSeq PE Cluster Kit v3-cBot-HS (Illumina) according to the manufacturer’s protocol. Finally, libraries were sequenced on an Illumina Novaseq 6000 platform, generating 150 bp paired-end reads.

### 2.5 Telomere tracking in live cells by dCas9 imaging

The *S. pyogenes* Cas9 gene, modified with D10A and H840A mutations (dCas9), was engineered to include a 2× SV40 nuclear localization sequence (NLS) and a 3× Flag-tag at the N-terminus, as well as GFP at the C-terminus ^15,16^. These dCas9 constructs were cloned into a lentiviral vector driven by the SFFV promoter. To establish a stable dCas9-expressing cell line, cells were transduced with the lentivirus, selected with 5 μg ml^-1^ puromycin for five days, and subsequently sorted via fluorescence-activated cell sorting (FACS) ^16,17^. The sgRNA plasmids were expressed under the U6 promoter, with the original plasmid backbone modified to include TagBFP as a transduction marker. The sgRNA sequences were amplified from an oligo template containing the 20-nucleotide telomere-targeting sequence: GUUAGGGUUAGGGUUAGGGUUA. For viral transduction, cells were incubated with a virus solution diluted threefold in DMEM medium supplemented with 10 μg ml^-1^ polybrene for 24 hours. For imaging telomere dynamics, fluorescence signals were captured using TIRF microscopy on a Nikon ECLIPSE Ti2 inverted microscope equipped with a Hamamatsu C11440-22C digital CMOS camera. The imaging parameters included an 80° incident angle, 200 ms intervals, and 300 ms exposure time. The laser power was set to 0.8 mW, and the gamma parameter was adjusted to 1^18,19^. At least 20 individual cells were examined for each case in the analysis. Telomere movement was tracked using CellProfiler software (version 4.2.25, https://cellprofiler.org/).

### 2.6 RNA-seq analysis

For trimming, trimming procedure was first applied to eliminate adapter sequences present in our data and to improve read quality from the FASTQ files. Adapter removal and quality trimming were carried out using Trimmomatic (v0.35). In all cases only reads with a Phred quality score > 20 and read length > 50 bp were selected for downstream analysis. Paired-end reads were removed if either read contained more than 10% ambiguous bases (N). Additionally, reads were discarded if more than 50% of their bases had a quality score of ≤ 5. Reads containing adapter sequences were also excluded from further analysis. Each sample produced approximately 80 million sequencing reads after quality control. We then use Salmon (v1.8.0) pseudoalignment to perform alignment, counting and normalization in one single step.

The isoform switching was analyzed using the IsoformSwitchAnalyzeR (v1.14.1) ^20^. The isoform switches were identified and quantified from RNA-seq data by using gtf annotation files from gencode (v47). IsoformSwitchAnalyzeR assesses isoform usage by calculating isoform fraction (IF) values, which represent the proportion of a gene’s total expression attributed to a specific isoform. This is determined by dividing the expression level of the isoform by the overall gene expression. Changes in isoform usage are quantified as the difference in isoform fraction (dIF), computed as IF2 − IF1. These dIF values are subsequently utilized to determine the isoform log2 fold change.

### 2.7 Statistical analysis

Each experiment was performed with a minimum of three replicates. All the analysis and comparisons were performed in R (v4.2.0). The DESeq2 (v1.46.0) package was utilized to analyze the differences between the doxycycline-treated groups and the untreated groups ^21^. Differential gene analysis was performed with a baseMean threshold of > 50 and a significance level of *p*-value < 0.05. Benjamini-Hochberg (BH) procedure was employed to control the false discovery rate (FDR) in multiple hypothesis testing (*q*-value). The resulting differential expressed gene (DEG) lists, ranked in descending order, were subsequently employed for Gene Ontology (GO) enrichment analysis using the clusterProfiler package (v3.20) ^22^. For significant levels of multiple group comparison, non-parametric Kruskal-Wallis test was used to compare the medians of multiple groups.

## 3. Results

### 3.1 Efficient knockdown of lamins and nesprins

The linker of nucleoskeleton and cytoskeleton (LINC) complex component nesprin-2 is a nuclear envelope protein that connects the nucleus to the cytoskeleton by interacting not only with actin filaments but also with microtubules through motor proteins such as dynein and kinesin. This structural linkage contributes to cellular architecture and facilitates mechanotransduction between the nuclear interior and the extracellular matrix (ECM) ^8,23^. lamin proteins weave a filamentous network in the inner nuclear interior to form lamina meshwork, which provides chromosomal anchoring sites to maintain genome organization. Nucleoplasmic lamins bind to chromatin and have been indicated to regulate chromatin accessibility and spatial chromatin organization ^24^. Lamins in the nuclear interior regulate gene expression by dynamically binding to heterochromatic and euchromatic regions, influencing epigenetic pathways and chromatin accessibility. They also contribute to chromatin organization and may mediate mechanosignaling ^25^. However, the contribution of nesprins and lamins to isoform switch and chromatin dynamics has not been fully understood ^7,10,26^.

We conducted RNA-seq and RT-qPCR to investigate how nesprins and lamins regulate gene expression following inducible shRNA-mediated knockdown. Our results demonstrate that lamin A affects RNA synthesis, *LMNA* alters chromatin conformation, and *SYNE2* modulates chromatin modification. *SYNE2* (nesprin-2) modulates chromatin modification (**Figure 1a**). In this work, shScramble, shLaminA, shLMNA, and shSYNE2 were denoted as comparisons between doxycycline-treated groups and untreated groups. Cells expressing doxycycline-inducible scramble shRNA were used as controls to rule out non-specific effects. To achieve gene knockdown, shRNAs targeting specific regions of lamin A, *LMNA*, and *SYNE2* transcripts were designed. The shRNA targeting the 3′ UTR of lamin A was used to selectively deplete lamin A (shLaminA), while a separate shRNA targeting the coding sequence (CDS) of *LMNA* was employed to reduce total *LMNA* expression (shLMNA). Additionally, the shRNA directed against the CDS of *SYNE2* was used to knock down nesprin-2 expression (shSYNE2). The targeting shRNA sequences were listed in **Supplementary Table S1**. The relative gene expression and log2 fold change were quantified in doxycycline-treated groups versus untreated groups for RT-qPCR and RNA-seq data, respectively. For RT-qPCR analysis in shScramble comparisons, the mean ± s.d. relative gene expression ratio was 0.96 ± 0.15 for lamin A, 0.95 ± 0.15 for *LMNA*, 1.03 ± 0.16 for *SYNE2*; For RNA-seq analysis in shScramble comparisons, the mean log2 fold change was 0.14 ± 0.10 for lamin A, 0.06 ± 0.05 for *LMNA*, 0.04 ± 0.11 for *SYNE2*. These data indicated that scramble shRNA did not change gene expression to the target genes. In contrast, targeting shRNAs significantly reduced the corresponding target genes. For RT-qPCR analysis, the mean ± s.d. relative gene expression ratio was 0.33 ± 0.01 for lamin A in shLaminA comparisons, 0.30 ± 0.02 for *LMNA* in shLMNA comparisons, 0.21 ± 0.04 for *SYNE2* in shSYNE2 comparisons; For RNA-seq analysis, the mean log2 fold change was −1.34 ± 0.08 for lamin A in shLaminA comparisons, 0.64 ± 0.06 for *LMNA* in shLMNA comparisons, −0.87 ± 0.22 for *SYNE2* in shSYNE2 comparisons (**Figure 1b**). Herein, we employed IsoformSwitchAnalyzeR to quantify log2 fold change of lamin A isoform (**Supplementary Figure S1**). Our findings demonstrated a significant reduction in the utilization of the lamin A isoform, while the expression levels of other *LMNA* gene isoforms remained unchanged. Together, our results indicated the efficient knockdown of lamin A, *LMNA* and *SYNE2* genes. Knockdown of lamin A specifically reduced the usage of its primary isoform, suggesting a potential role in chromatin architecture regulation, while other *LMNA* isoforms remained unaffected, highlighting a selective effect. Together, our results indicated the efficient knockdown of lamin A, *LMNA*, and *SYNE2* genes.

**Figure 1.**
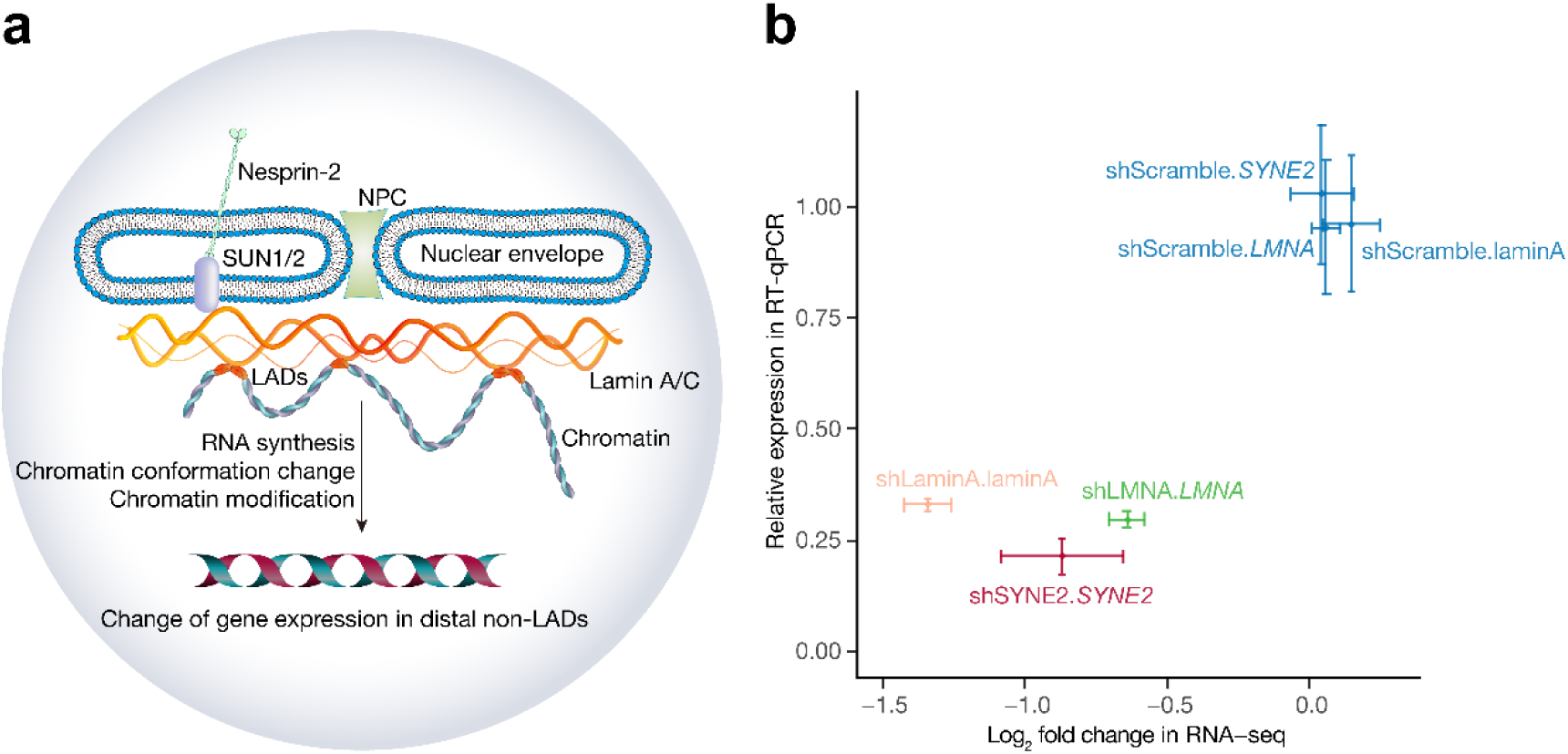
Knockdown of lamins and nesprins alters gene expression. **a)** Schematic illustration of the regulation of gene expression by lamins and nesprins. The lamins and nesprins could affect gene expression in distal non-LADs through altering RNA synthesis, chromatin conformation change, and chromatin modification. NPC, nuclear pore complex; LADs, lamina-associated domains. **b)** Knockdown of lamin A, *LMNA*, and *SYNE2* mRNA levels by targeting shRNAs (shLaminA, yellow; shLMNA, green; shSYNE2, red) was quantified by RT-qPCR and RNA-seq, respectively. In contrast, the relative gene levels of targeting genes (lamin A, *LMNA*, *SYNE2*) are unchanged in the U2OS cells with scramble shRNA construct (shScramble, blue). U2OS cells were treated or untreated with 100 ng ml^-1^ doxycycline for 48 h. The isoform log2 fold change of lamin A was calculated by IsoformSwitchAnalyzeR package. Data represent mean ± s.d. RT-qPCR, n = 7 per group, *p*-value < 0.05; RNA-seq, n = 3 per group, *q*-value < 0.05.

### 3.2 Impact of lamins and nesprins on RNA biosynthesis, chromatin conformation, and modification

Correlation heatmap and principal component analysis (PCA) plot revealed distinct gene expression profiles across the shScramble, shLaminA, shLMNA, and shSYNE2 groups (**Supplementary Figure S2**). Next, by performing DE analysis with DESeq2, our data showed that depletion of lamins and nesprins could significantly lead to the alteration of gene expression, including the protein-coding mRNAs and long non-coding lncRNAs (**Supplementary Table S3**). To examine pathway enrichment by the DEGs in the shLaminA, shLMNA, and shSYNE2 comparisons. GO analysis on biological process (BP) was conducted by clusterProfiler package, which was based on the hypergeometric test of descending ranking of log2 fold change of DEGs. No significant BP pathway was enriched for shScramble comparison (**Supplementary Figure S3**). Intriguingly, RNA synthesis pathways were enriched in the knockdown of lamin A isoform, including purine-containing compound metabolic process and ribose phosphate metabolic process, which ensure the synthesis of the ribonucleotide triphosphates (NTPs) that required for RNA synthesis (**Figure 2a**). In contrast, the knockdown of the *LMNA* gene was associated with an enrichment of DNA conformation change, suggesting that *LMNA* may regulate alterations in the spatial organization of chromatin (**Figure 2b**). In addition, the knockdown of the *SYNE2* gene was linked to the ncRNA metabolic process and covalent chromatin modification, indicating that the impact of *SYNE2* in the regulation of chromatin accessibility, remodeling and gene expression (**Figure 2c**).

**Figure 2.**
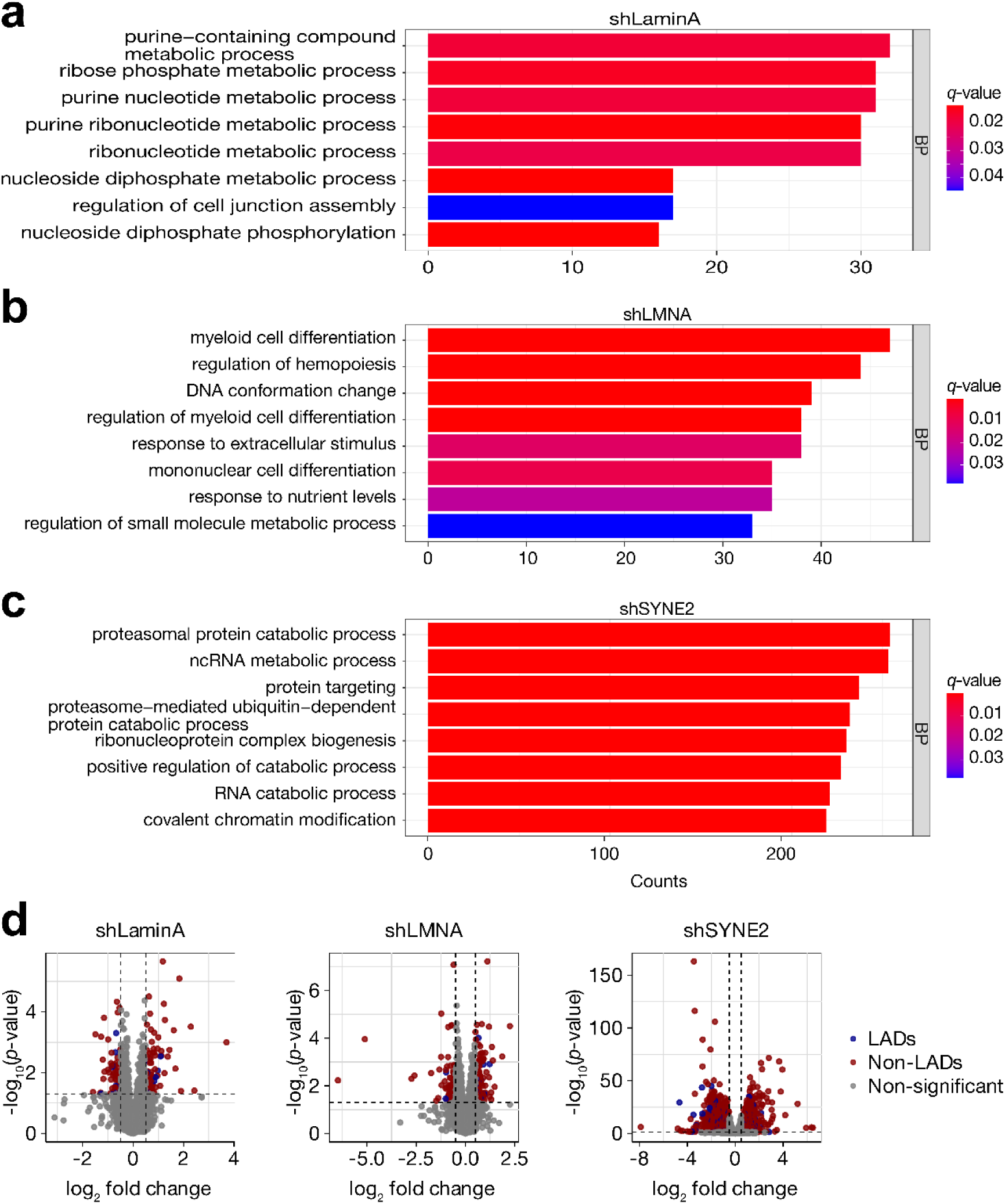
Impact of lamins and nesprins on RNA biosynthesis, chromatin conformation, and modification. DEGs were identified using the DESeq2 package by comparing the doxycycline-treated group to the untreated group. Gene Ontology (GO) enrichment analysis of DEGs was conducted using the clusterProfiler package. Gene counts are displayed on the *x*-axis. The top pathways were selected based on a *q*-value cutoff of < 0.05 for shLaminA (**a**), shLMNA (**b**), and shSYNE2 (**c**), respectively. **d**) Volcano plots illustrating the distribution of DEGs located in lamina-associated domains (LADs) and non-LAD regions. DEGs were defined by an absolute log₂ fold change > 0.5 and *p*-value < 0.05.

Furthermore, differential expression analysis revealed that the majority of DEGs following depletion of lamins and nesprins were located outside lamina-associated domains (non-LADs). Specifically, for shLaminA knockdown, 8 DEGs within LADs were downregulated and 8 were upregulated, whereas 59 non-LAD DEGs were downregulated and 79 were upregulated. For shLMNA, 7 LAD-associated DEGs were downregulated and 15 were upregulated, with 88 downregulated and 140 upregulated DEGs in non-LAD regions. In the case of shSYNE2 knockdown, 161 LAD DEGs were downregulated and 108 were upregulated, while 2,009 non-LAD DEGs were downregulated and 1,851 were upregulated (**Figure 2d, Supplementary Table S4**). These results indicate that the transcriptional changes resulting from the loss of lamins or nesprins predominantly occur at non-LAD genomic regions.

The percentage of DEGs was consistently higher in non-LADs, which are gene rich and transcriptionally active, whereas LADs, known to be enriched for silent or lowly expressed genes, showed fewer expression changes. These findings are consistent with previous studies demonstrating that active genes are more prevalent in non-LADs and that LAD associated genes are generally repressed or less responsive to perturbation ^27,28^. Together, these results support a model in which lamins and nesprins influence gene expression through both structural organization and promoter proximal interactions, particularly within euchromatic nuclear regions ^10,26,29^.

### 3.3 Role of lamins and nesprins in isoform switches

To uncover transcript-specific regulatory changes, we performed isoform-level differential expression analysis. Many genes produce functionally distinct isoforms, and shifts in their usage can occur without changes in total gene expression, making isoform-level analysis essential for detecting subtle but meaningful transcriptional regulation. To investigate how lamins and nesprins would influence isoform switches, we performed isoform switches analysis by IsoformSwitchAnalyzeR package. To identify key isoform changes upon *LMNA* knockdown, we ranked genes by ascending *q*-values of isoform switching. The top ten genes with significant isoform alterations included *TARS1*, *PPP1R13L*, *ATP6AP2*, *NBL1*, *JADE1*, *AGPS*, *Lnc-ZFAT-1*, *RBM11*, and *PEX19* (**Figure 3a, Supplementary Table S5**). These genes are functionally involved in diverse processes such as protein synthesis (*TARS1*), chromatin remodeling (*JADE1*), RNA processing (*RBM11*), and protein degradation (*ATP6AP2*, *PEX19*).

**Figure 3.**
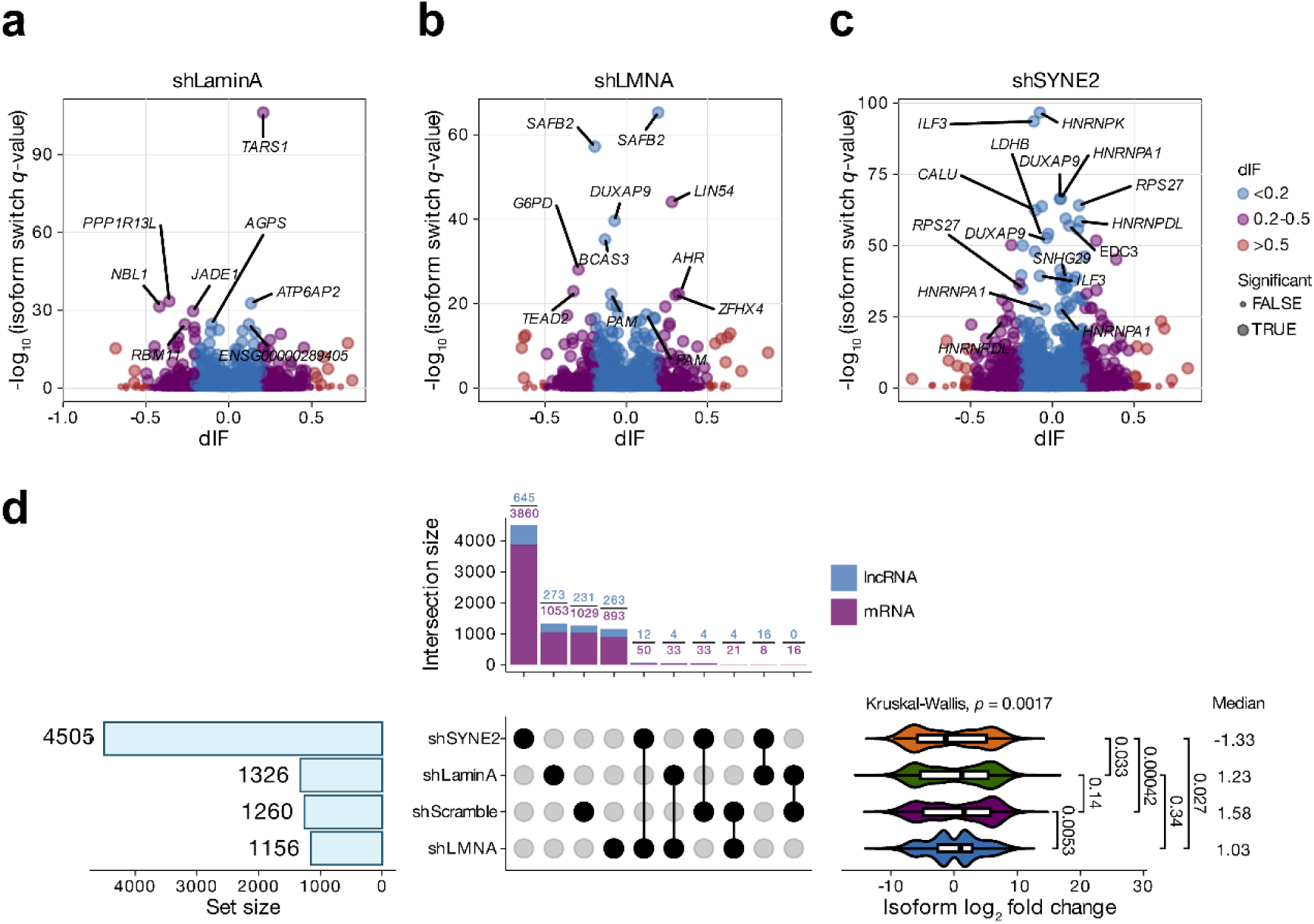
Roles of lamins and nesprins in isoform switching. Isoform switching analysis reveals differential expression of alternative transcript variants between conditions, highlighting a shift in predominant isoform usage. **a, b, c)** Volcano plots illustrating the dIF of differentially expressed isoform transcripts against the −log10 of the isoform switch *q*-values in shLaminA (**a**), shLMNA (**b**), and shSYNE2 (**c**). Isoform switching was detected using IsoformSwitchAnalyzeR (n = 3 per group, cutoff dIF > 0.1, *q*-value < 0.05). **a**) Lamin A knockdown leads to isoform switching in chromatin remodeling (*JADE1*) and alternative splicing regulation (*RBM11*). **b**) *LMNA* knockdown results in isoform switching of transcriptional repressors (*SAFB2*, *LIN54*) and transcription factors (*TEAD2*, *AHR*). **c**) *SYNE2* knockdown significantly alters isoform ratios of RNA-binding proteins (*HNRNPK*, *ILF3*, *HNRNPA1*). **d**) UpSet plot displaying the number and overlap of differentially expressed isoforms in protein-coding mRNAs and lncRNAs, as well as the log2 fold change of isoform transcripts in shLaminA, shLMNA, and shSYNE2 conditions. Intersection sizes indicate the number of shared DEGs between knockdowns. Statistical analysis was performed using the non-parametric Kruskal-Wallis test to compare medians across multiple groups.

A separate set of top-ranking isoform switch events in LMNA knockdown included *SAFB2*, *LIN54*, *DUXAP9*, *BCAS3*, *G6PD*, *TEAD2*, *AHR*, *PAM*, and *ZFHX4* (**Figure 3b**). These genes are enriched in functions related to transcriptional repression (*SAFB2*, *LIN54*), transcription factor activity (*TEAD2*, *AHR*, *ZFHX4*), and post-translational modification (*PAM*).

In another subset of significantly affected genes (**Figure 3c**), the top isoform switches included *HNRNPK*, *ILF3*, *HNRNPA1*, *DUXAP9*, *RPS27*, *LDHB*, *CALU*, *SNHG29*, *HNRNPDL*, and *EDC3*. These genes are implicated in RNA splicing and stability (*HNRNPK*, *ILF3*, *HNRNPA1*, *HNRNPDL*), mRNA decay (*EDC3*), and protein synthesis or folding (*RPS27*, *LDHB*).

Nesprin-2 depletion induced a substantially larger number of DE isoforms (n = 4505) compared to lamin A (n = 1326) and *LMNA* (n = 1156) gene knockdown. This included 645 DE long non-coding lncRNAs and 3860 DE mRNAs in nesprin-2 knockdown cells, as well as 273 DE lncRNAs and 1053 DE mRNAs following lamin A silencing, and 263 DE lncRNAs and 893 DE mRNAs after *LMNA* reduction (**Figure 3d**). While the number of DEGs identified for lamins was limited, a significant number of isoform switches was observed following lamin knockdown. Interestingly, the UpSet plot analysis revealed a high degree of specificity in the gene expression changes induced by each gene knockdown. Minimal overlaps were observed between the differentially expressed genes in shSYNE2, shLaminA, and shLMNA conditions: shSYNE2 and shLaminA (16 lncRNAs, 8 mRNAs), shSYNE2 and shLMNA (12 lncRNAs, 50 mRNAs), and shLaminA and shLMNA (4 lncRNAs, 33 mRNAs), suggesting distinct downstream effects of each gene knockdown. Relative to the shScramble control (median isoform log2 fold change, 1.58), knockdown of *LMNA*, lamin A, and *SYNE2* resulted in substantial changes in isoform expression, with median log2 fold changes of 1.03, 1.23, and −1.33, respectively. Notably, shSYNE2 induced overall reduction of isoform switches. Collectively, our results demonstrated that knockdown of each gene, including the lamin A isoform, *LMNA*, and *SYNE2*, led to distinct and highly specific changes in isoform switching. Our analysis demonstrated that depletion of lamins and nesprins induced significant alterations in specific transcript isoforms, indicating regulatory changes in alternative splicing or transcription initiation that are not captured by gene-level differential expression analysis.

### 3.4 Regulation of gene expression in distal non-LAD regions by lamins and nesprins

The majority of genes located within LADs tend to be transcriptionally repressed or expressed at low levels. This is because LADs are associated with heterochromatin, a tightly packed form of DNA that is generally inaccessible to the cellular machinery required for gene expression ^14,30^. Lamin mutations and levels have been shown to disrupt LAD organization and gene expression that have been implicated in various diseases, including cancer and laminopathies ^31,32^. Next, we investigated the chromosome localizations of DE mRNAs in the genome-wide. LADs exhibit dynamic reorganization and changes in gene expression during cellular differentiation ^28^. Although genes within LADs are generally transcriptionally silent or expressed at low levels ^33^, some LAD-resident genes remain active and can be transcriptionally modulated in response to specific stimuli, such as T cell activation ^34^. The Venn diagram analysis revealed limited overlap between DEGs resulting from knockdown of lamin A (shLaminA), *LMNA* (shLMNA), or *SYNE2* (shSYNE2) and genes located within lamina-associated domains (LADs). Specifically, only a small subset of DEGs intersected with LAD-associated genes across all three knockdowns, suggesting that the majority of transcriptional changes occur outside LAD regions (**Figure 4a**). Consistent with prior studies, depletion of lamin A, *LMNA*, and *SYNE2* did not significantly impact LAD-associated genes, reinforcing the idea that their primary function in gene regulation may occur in distal non-LAD chromatin regions ^14^.

**Figure 4.**
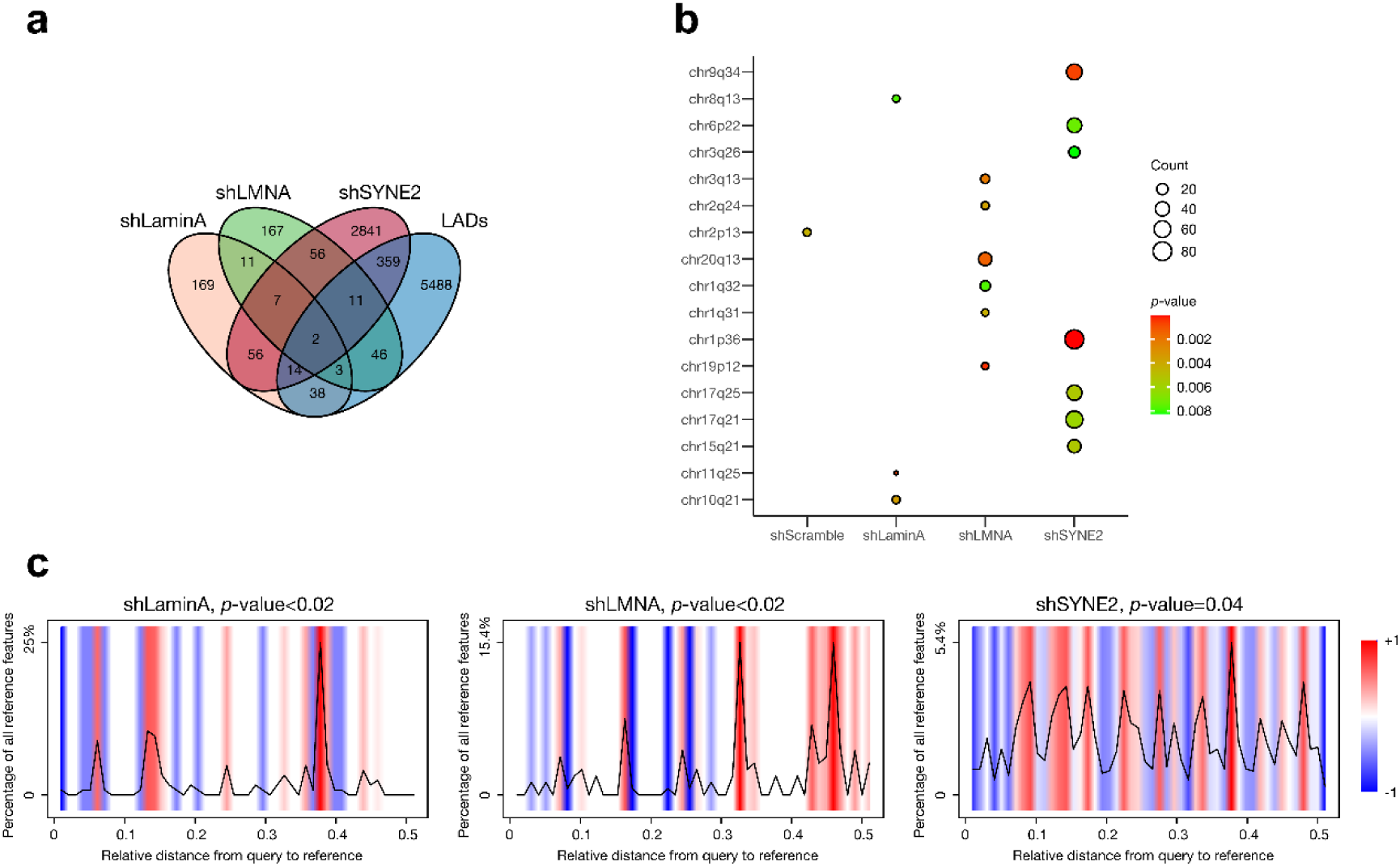
Regulation of gene expression in distal non-LAD regions by lamins and nesprins. **a)** Venn diagram illustrating the number of DE mRNAs in shLaminA (yellow), shLMNA (green), and shSYNE2 (red), with minimal overlap with genes located in LADs (blue). LAD-associated genes were identified using a BED file derived from lamin A/C ChIP-seq data ^54^. DEGs were identified using thresholds of absolute log2 fold change > 0.5 and *p*-value < 0.05. **b)** GO enrichment analysis showing the genomic distribution of DEGs across chromosome arms. for shLaminA, shLMNA, and shSYNE2. The *p*-values (blue to red), and the counts of DE genes mapped to the reference chromosome coordinates are shown. Depletion of lamin A affects chr10q12, *LMNA* affects chr20q13, and *SYNE2* affects chr9q34, chr1q36. **c)** DE genes from shLaminA, shLMNA, and shSYNE2 are spatially distant from LADs. GenometriCorr analysis (v1.1.24) was used to assess the spatial relationship between DEGs (query) and LADs (reference) ^55^. The *x*-axis shows the relative genomic distance between each DEG and the nearest LAD, scaled from –1 (far upstream) to 1 (far downstream), with 0 indicating closest proximity. The *y*-axis represents the density of DEGs at each distance bin. A color gradient indicates deviation from a randomized null distribution: red signifies enrichment (closer than expected), and blue signifies depletion. Statistical significance was determined using the Jaccard test (*p* < 0.05).

Interestingly, the DEGs from depletion of lamin A, *LMNA*, and *SYNE2* were dispersed in different chromosome loci when mapping genes to human hg19 genome reference. The DEGs from depletion of lamin A were located in chr8q13, chr10q21, and chr11q25. The DEGs from depletion of *LMNA* were mainly located in chr20q13, chr3q13, and chr19p12. The DEGs from depletion of *SYNE2* were mainly located in chr9q34, chr1p36, and chr17q21 (**Figure 4b**). To confirm that the DEGs were not located in LADs, we measured the spatial distance between LADs and DEGs in shLaminA, shLMNA, and shSYNE2 through GenometriCorr package. Our data suggested that those DEGs are spatially distant from LADs (**Figure 4c**). Together, our results show that depletion of lamin A, *LMNA*, and *SYNE2* altered gene expression in distal non-LAD regions.

### 3.5 Modulation of lncRNA-mRNA interactions in the nucleus by lamins and nesprins

Consistent with published results, our data showed lower lncRNA expression compared to mRNA (**Supplementary Figure S4**) ^35^. Interestingly, mRNA expression showed a unimodal distribution in the log2 transcript per million (Log2TPM) density plot. In contrast, the lncRNA expression profiling demonstrated multimodal distribution with multiple peaks (**Figure 5a**). In addition, analysis using DeepLncLoc predicted that lncRNAs are primarily located in the nucleus (**Figure 5b**) ^36^. These results suggest that mRNA-lncRNA interactions primarily occur in the nucleus.

**Figure 5.**
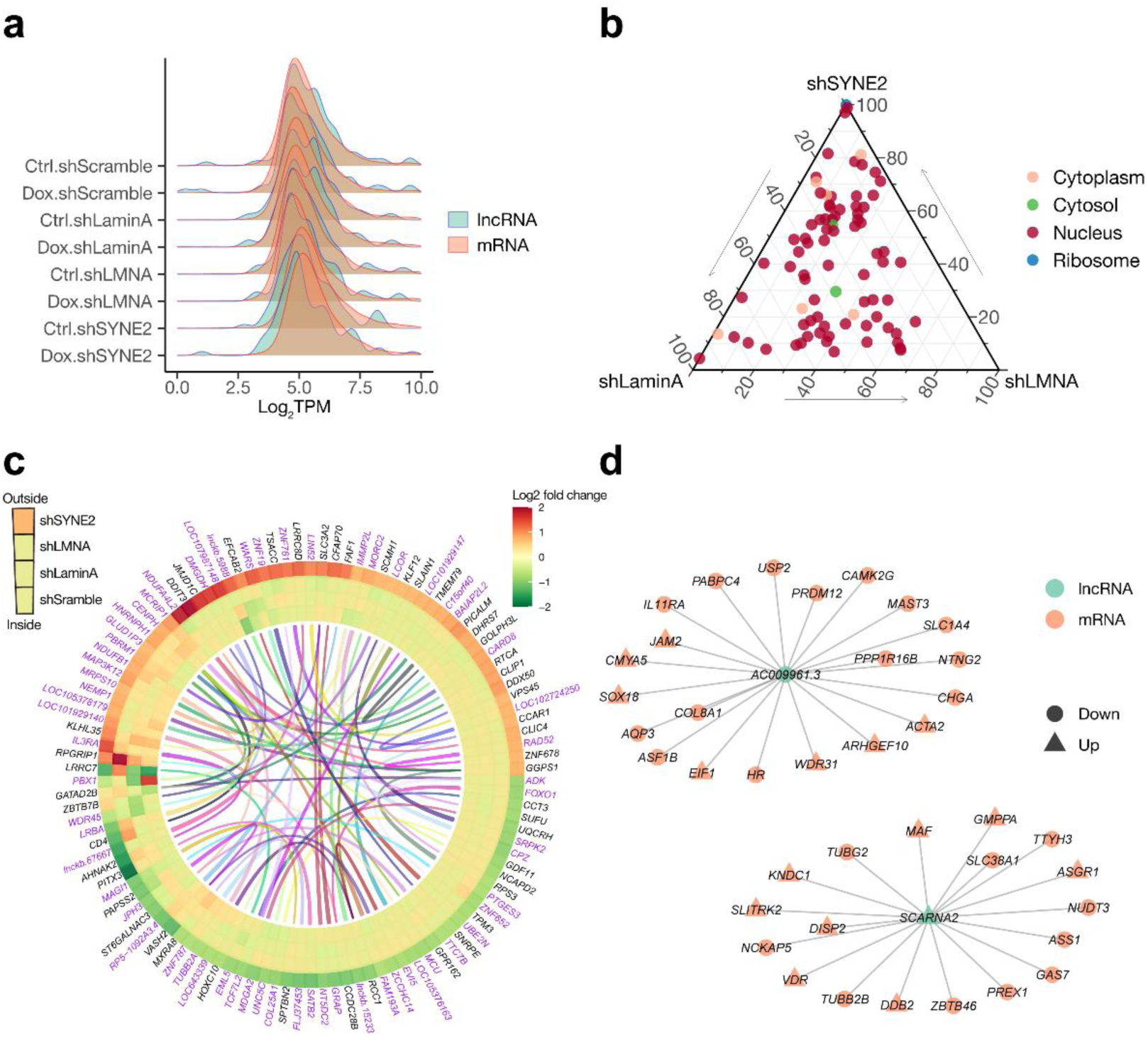
Modulation of lncRNA-mRNA Interactions by Lamins and Nesprins. **a)** Expression distribution of lncRNAs versus mRNAs in log2 transcript per million (Log2TPM), highlighting differences in transcript abundance. **b)** DeepLncLoc predictions showing nuclear localization of identified lncRNAs. **c)** Cis-acting lncRNA-mRNA interaction network predicted based on highly correlated expression levels within 10 kb windows (Pearson correlation > 0.8, *q*-value < 0.05). **d)** Coexpression network analysis of lncRNA-mRNA interactions. Nodes represent transcripts; edges denote coexpression relationships (Pearson correlation > 0.8, *q*-value < 0.05).

Furthermore, We identified cis-acting mRNA-lncRNA pairs by analyzing highly correlated expression within 10 kb genomic windows (**Figure 5c**). Additionally, the lncTar algorithm and Pearson correlation test were employed to analyze lncRNA-mRNA coexpression networks, enabling the identification and validation of trans-acting and/or cis-acting lncRNA-mRNA interaction networks ^37^. We identified two interaction networks between two lncRNAs (green nodes) and their associated mRNAs (orange nodes). Upregulated genes are represented as triangles, while downregulated genes are shown as circles. Two clusters are centered around the lncRNAs *AC009961.3* and *SCARNA2*, highlighting their regulatory roles in specific gene expression pathways. The networks underscored the potential mechanistic influence of lncRNAs in cellular processes and their functional impact on mRNA expression (**Figure 5d**).

### 3.6 Increased telomere dynamics with lamins and nesprins depletion

Although telomeres are not uniformly tethered to the nuclear lamina, they can transiently associate with the nuclear periphery, particularly during post-mitotic nuclear reassembly, through interactions involving SUN1 and RAP1 ^38^. Given that lamins and nesprins are key components of the nuclear envelope that regulate chromatin organization and mechanics ^39,40^, we examined telomere dynamics as a proxy for changes in nuclear architecture. Using EGFP-tagged dCas9 to label telomeric regions in live U2OS cells, we assessed whether knockdown of these proteins leads to increased telomere mobility, reflecting a loss of structural constraint or altered chromatin–nuclear envelope interactions ^17^. Representative fluorescence images of telomeres labeled with EGFP-dCas9 in U2OS cells are shown in **Figure 6a**. Notably, depletion of lamin A, *LMNA*, and *SYNE2* resulted in significantly increased genome dynamics and a larger nuclear area scanned. This effect was quantified through telomere mean square displacement (MSD) analysis and the measurement of traveled distance within a 30-second imaging window (**Figure 6b, c and Supplementary Movie S1**). These findings support the concept that lamins play a critical role in regulating chromatin dynamics ^10^. Collectively, our results highlight a regulatory function of lamins and nesprins in controlling chromatin behavior.

**Figure 6.**
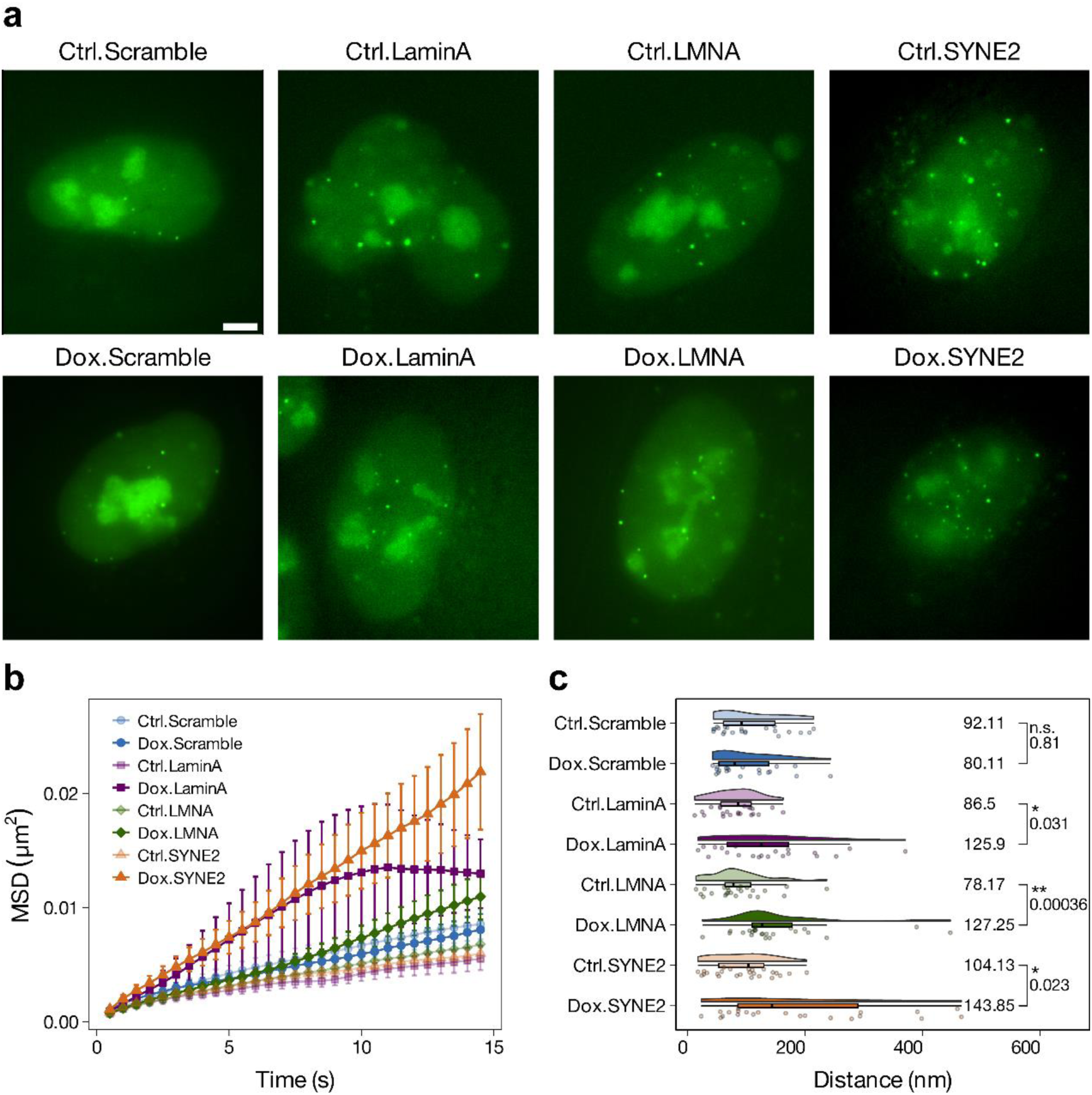
Increased telomere dynamics following depletion of lamin A, LMNA, and SYNE2. **a)** Representative fluorescence images of telomeres labeled with EGFP-dCas9 in U2OS cells. Cells expressing dCas9-EGFP, targeted to telomeric repeats, were imaged via TIRF microscopy. Scale bar = 5 µm. **b)** MSD analysis of telomere mobility over 30 s at 500 ms intervals in control and knockdown conditions. MSD curves represent n > 20 cells per group. **c)** Total telomere movement range tracked over a 30 s imaging window. Annotated numbers indicate the median values for each group. Statistical analysis was performed using the non-parametric Kruskal-Wallis test, with a significance threshold of \**p*-value < 0.05, \*\**p*-value < 0.001, and n.s. *p*-value > 0.05.

To probe how nuclear envelope components regulate chromatin dynamics, we tracked telomeres as a representative genomic locus whose mobility reflects changes in nuclear mechanics and chromatin organization. Although telomeres are not stably tethered to the nuclear lamina, their motion can be influenced by nuclear architecture and transient peripheral associations ^38^. Upon depletion of lamin A, *LMNA*, or *SYNE2*, we observed significantly increased telomere mobility and nuclear area explored, quantified by mean square displacement and net displacement (**Figure 6b–c, Supplementary Movie S1**). These changes likely reflect altered chromatin–lamina interactions or disrupted nuclear mechanical constraints, consistent with prior studies showing that lamins modulate chromatin dynamics and nuclear stiffness ^39,40,41^. Thus, our findings support a role for lamins and nesprins in constraining chromatin motion through nuclear structural integrity.

## 4. Discussion

LADs occupy roughly 40% of the genome, although these regions are not uniformly bound to the nuclear lamina in every cell within a population. These domains are defined by their physical association with the nuclear lamina and are distinguished by a low gene density, typically containing about two to three genes per megabase. This is considerably lower than the average gene density of the human genome, which contains approximately eight genes per megabase. These regions are enriched with repressive chromatin features, limiting transcriptional activity compared to non-LAD regions, which are typically more transcriptionally active ^13,42^.

Beyond their well-established role in tethering heterochromatin at the nuclear periphery through lamina-associated domains (LADs), A-type lamins (lamins A and C) also localize to the nuclear interior, where they contribute to chromatin organization and gene regulation independently of LADs ^40,43^. Nuclear lamins can form intranuclear foci that associate with active chromatin and are implicated in supporting transcriptional activity. Additionally, both lamins and nesprins participate in diverse protein-protein interactions that may influence transcriptional regulation. For example, lamin A/C interacts with the retinoblastoma protein (Rb) to modulate E2F-dependent transcription ^44^, and with c-Fos to regulate its nuclear retention and activity ^45^. While β-catenin acts as a co-activator in Wnt signaling relies on nuclear translocation and interaction with transcriptional complexes, and evidence suggests that nuclear architecture and envelope components, including nesprins, can influence this process ^46^. Therefore, the observed gene expression changes following depletion of lamins or nesprins are likely not restricted to genes located within lamina-associated domains (LADs), but may also result from broader perturbations in nuclear architecture and transcriptional regulatory networks. This is consistent with our findings that lamins and nesprins influence gene expression in distal, non-LAD regions.

Our findings reveal distinct functional roles for Lamin A, *LMNA*, and *SYNE2* in chromatin regulation and gene expression, providing key insights into the spatial organization of the genome. Notably, lamin A depletion led to enrichment of pathways associated with RNA biosynthesis, supporting its previously suggested role in transcriptional activation and ribonucleotide metabolism ^47^. This aligns with prior studies indicating that lamin A contributes to chromatin accessibility and RNA polymerase activity ^48^. These findings further underscore the functional relevance of lamin A in coordinating transcriptional programs through modulation of nuclear architecture. In contrast, *LMNA* knockdown led to differential expression of genes enriched in pathways related to chromatin organization, suggesting potential disruptions in chromatin regulatory networks. Although direct measurements of chromatin conformation were not performed, these transcriptional changes indicate that *LMNA* may contribute to maintaining nuclear architecture and genomic stability, which aligns with its established involvement in laminopathies and genome integrity disorders. The findings that DEGs are predominantly located in non-LAD regions highlight a unique regulatory aspect of lamins and nesprins, emphasizing their spatial specificity in gene expression. Moreover, the modulation of lncRNA-mRNA interactions by these proteins unveils a layered regulatory network, further illustrating the nuclear envelope’s influence on transcriptional dynamics. The isoform switching patterns observed following knockdown of lamin A and *LMNA* provide further insight into transcriptomic flexibility. Our data show that knockdown with shLaminA, which specifically targets the 3’ UTR unique to the lamin A isoform, selectively reduced lamin A expression without affecting other *LMNA* isoforms. Given that lamin isoforms are known to differentially impact nuclear stiffness and mechanotransduction, these findings raise the possibility that specific lamin isoforms may be preferentially required for distinct chromatin interactions and nuclear functions.

These findings align with studies showing that nuclear envelope dysfunction, as seen in laminopathies and cancer, leads to altered nuclear mechanics and chromatin misregulation ^4,49,50^, where altered nuclear mechanics and chromatin misregulation are hallmarks of disease progression. The observed enhancement of chromatin dynamics upon depletion of lamin A, *LMNA*, and *SYNE2*, as visualized through dCas9-based live chromosome imaging, suggests that these nuclear envelope proteins contribute to chromatin rigidity and nuclear compartmentalization. Their depletion leads to increased chromatin fluidity, which may underlie genomic instability and misregulated transcription in disease states.

Our findings reinforce the pivotal roles of nuclear envelope proteins lamin A, LMNA and nesprin 2 in regulating chromatin organization, chromatin mobility, and gene expression. These results are consistent with and extend prior studies investigating the consequences of lamin depletion. For instance, increased chromatin mobility following the loss of lamin A/C has been previously demonstrated using live-cell imaging approaches ^10,11^, supporting our observations of nuclear structural relaxation and chromatin redistribution. Additionally, proteomic profiling following lamin A depletion has been extensively documented across both cellular and mouse models, providing valuable insights into the molecular consequences of nuclear envelope disruption ^32,51^. While these earlier studies provide a strong foundation, our work contributes novel insights by integrating isoform-specific perturbations with spatial chromatin measurements. This approach emphasizes context-dependent regulatory mechanisms that involve not only lamina-associated regions but also nesprin-associated domains and distal genomic loci, thereby expanding the current understanding of nuclear envelope protein function in gene regulation.

Our findings clarify how chromatin behavior is regulated, advancing the broader understanding of nuclear architecture’s role in cellular homeostasis. These insights may have broad implications for understanding nuclear envelope-associated diseases, particularly those involving mutations in *LMNA* or *SYNE2*, and may help guide future therapeutic strategies targeting nuclear envelope dysfunction.

Interestingly, although both shLMNA and shLaminA constructs target lamin A, with shLMNA additionally depleting lamin C, the DEGs identified under these two conditions show limited overlap. This unexpected finding suggests that depletion of lamin C in the shLMNA condition may trigger distinct or compensatory transcriptional responses that are not elicited by lamin A knockdown alone. Furthermore, variation in shRNA efficiency or off-target effects may contribute to these differences. Notably, despite directly targeting LMNA, the overlap in DEGs between the two conditions remained limited under our stringent threshold criteria. Together, these observations highlight the complex and non-linear regulatory roles of lamin isoforms in gene expression and underscore the need for further mechanistic studies to dissect their individual and combined contributions ^52,53^.

## 5. Conclusion

In summary, this study demonstrates the intricate roles of lamin A, *LMNA*, and *SYNE2* in regulating distal chromatin and gene expression. Inducible knockdowns and transcriptomic analysis revealed their distinct roles in RNA synthesis, chromatin conformation, and modification. Using dCas9-based live chromosome imaging, enhanced chromatin dynamics were observed for the depletion of lamin A, *LMNA*, and *SYNE2* genes. These findings deepen our understanding of nuclear envelope-linked gene regulation and its role in chromatin-related diseases. Future research should explore the broader impact of these mechanisms in diverse cellular contexts, advancing potential therapeutic strategies targeting nuclear envelope dysfunction.

## 6. Data availability

BED file was retrieved from lamin A/C CHIP-seq (https://github.com/kohta-ikegami/pS22-LMNA/blob/master/E1_BJ5ta_LAD.bed)^54^. The raw count data generated by Salmon are provided in Supplementary Table S6. All raw and processed data are available in the Supplementary Tables. All raw sequencing data produced in this study have been deposited in the NCBI SRA database under accession number PRJNA1212085.

## Compliance and ethics

The authors declare no conflicts of interest or personal relationships influencing this work.

## Acknowledgements

This work is partially supported by the National Natural Science Foundation of China (Grant No. 32350410397); Shenzhen Medical Research Funds (Grant No. D2301002); and the Science, Technology, and Innovation Commission of Shenzhen Municipality (Grant Nos. JCYJ20240813112016022, JCYJ20220530143014032, JCYJ20230807113017035, KCXFZ20211020163813019); Tsinghua Shenzhen International Graduate School Cross-disciplinary Research and Innovation Fund Research Plan, JC2022009; and Bureau of Planning, Land and Resources of Shenzhen Municipality (2022) 207.

## Notes

### Competing Interest Statement

The authors have declared no competing interest.

### Summary of Updates

This version has been substantially revised. We clarified the specific roles of lamin A, LMNA, and SYNE2 in RNA synthesis, chromatin organization, and chromatin modification, respectively. The Discussion and Conclusion sections were extensively rewritten to integrate these findings and clarify the distinct roles of nuclear envelope proteins in chromatin dynamics.

## References

1. Buchwalter, A., Kaneshiro, J. M. & Hetzer, M. W. Coaching from the sidelines: the nuclear periphery in genome regulation. Nat Rev Genet 20, 39–50 (2019).

2. Karoutas, A. & Akhtar, A. Functional mechanisms and abnormalities of the nuclear lamina. Nat Cell Biol 23, 116–126 (2021).

3. Irianto, J., Pfeifer, C. R., Ivanovska, I. L., Swift, J. & Discher, D. E. Nuclear Lamins in Cancer. Cel. Mol. Bioeng. 9, 258–267 (2016).

4. Bell, E. S. et al. Low lamin A levels enhance confined cell migration and metastatic capacity in breast cancer. Oncogene 41, 4211–4230 (2022).

5. Scaffidi, P., Gordon, L. & Misteli, T. The Cell Nucleus and Aging: Tantalizing Clues and Hopeful Promises. PLOS Biology 3, e395 (2005).

6. Shah, P. P. et al. Pathogenic LMNA variants disrupt cardiac lamina-chromatin interactions and de-repress alternative fate genes. Cell Stem Cell 28, 938–954.e9 (2021).

7. Swift, J. et al. Nuclear Lamin-A Scales with Tissue Stiffness and Enhances Matrix-Directed Differentiation. Science 341, 1240104 (2013).

8. Rajgor, D. & Shanahan, C. M. Nesprins: from the nuclear envelope and beyond. Expert Reviews in Molecular Medicine 15, e5 (2013).

9. Li Mow Chee, F. et al. Mena regulates nesprin-2 to control actin–nuclear lamina associations, trans-nuclear membrane signalling and gene expression. Nat Commun 14, 1602 (2023).

10. Bronshtein, I. et al. Loss of lamin A function increases chromatin dynamics in the nuclear interior. Nat Commun 6, 8044 (2015).

11. Ranade, D., Pradhan, R., Jayakrishnan, M., Hegde, S. & Sengupta, K. Lamin A/C and Emerin depletion impacts chromatin organization and dynamics in the interphase nucleus. BMC Molecular and Cell Biology 20, 11 (2019).

12. van Steensel, B. & Furlong, E. E. M. The role of transcription in shaping the spatial organization of the genome. Nat Rev Mol Cell Biol 20, 327–337 (2019).

13. Briand, N. & Collas, P. Lamina-associated domains: peripheral matters and internal affairs. Genome Biology 21, 85 (2020).

14. van Steensel, B. & Belmont, A. S. Lamina-Associated Domains: Links with Chromosome Architecture, Heterochromatin, and Gene Repression. Cell 169, 780– 791 (2017).

15. Bandaria, J. N., Qin, P., Berk, V., Chu, S. & Yildiz, A. Shelterin Protects Chromosome Ends by Compacting Telomeric Chromatin. Cell 164, 735–746 (2016).

16. Qin, P. et al. Live cell imaging of low- and non-repetitive chromosome loci using CRISPR-Cas9. Nat Commun 8, 14725 (2017).

17. Chen, B. et al. Dynamic Imaging of Genomic Loci in Living Human Cells by an Optimized CRISPR/Cas System. Cell 155, 1479–1491 (2013).

18. Lei, Z. et al. Detection of Frog Virus 3 by Integrating RPA-CRISPR/Cas12a-SPM with Deep Learning. ACS Omega 8, 32555–32564 (2023).

19. He, Q. et al. Unraveling the influence of CRISPR/Cas13a reaction components on enhancing trans-cleavage activity for ultrasensitive on-chip RNA detection. Microchim Acta 191, 466 (2024).

20. Vitting-Seerup, K. & Sandelin, A. IsoformSwitchAnalyzeR: analysis of changes in genome-wide patterns of alternative splicing and its functional consequences. Bioinformatics 35, 4469–4471 (2019).

21. Love, M. I., Huber, W. & Anders, S. Moderated estimation of fold change and dispersion for RNA-seq data with DESeq2. Genome Biology 15, 550 (2014).

22. Wu, T. et al. clusterProfiler 4.0: A universal enrichment tool for interpreting omics data. The Innovation 2, 100141 (2021).

23. Luxton, G. G. & Starr, D. A. KASHing up with the nucleus: novel functional roles of KASH proteins at the cytoplasmic surface of the nucleus. Current Opinion in Cell Biology 28, 69–75 (2014).

24. Borsos, M. et al. Genome–lamina interactions are established de novo in the early mouse embryo. Nature 569, 729–733 (2019).

25. Gruenbaum, Y. & Foisner, R. Lamins: Nuclear Intermediate Filament Proteins with Fundamental Functions in Nuclear Mechanics and Genome Regulation. Annual Review of Biochemistry 84, 131–164 (2015).

26. Wang, Y. et al. Lamin A/C-dependent chromatin architecture safeguards naïve pluripotency to prevent aberrant cardiovascular cell fate and function. Nat Commun 13, 6663 (2022).

27. Guelen, L. et al. Domain organization of human chromosomes revealed by mapping of nuclear lamina interactions. Nature 453, 948–951 (2008).

28. Peric-Hupkes, D. et al. Molecular Maps of the Reorganization of Genome-Nuclear Lamina Interactions during Differentiation. Molecular Cell 38, 603–613 (2010).

29. Naetar, N., Ferraioli, S. & Foisner, R. Lamins in the nuclear interior − life outside the lamina. Journal of Cell Science 130, 2087–2096 (2017).

30. Guelen, L. et al. Domain organization of human chromosomes revealed by mapping of nuclear lamina interactions. Nature 453, 948–951 (2008).

31. Perovanovic, J. et al. Laminopathies disrupt epigenomic developmental programs and cell fate. Sci Transl Med 8, 335ra58 (2016).

32. Bell, E. S. et al. Low lamin A levels enhance confined cell migration and metastatic capacity in breast cancer. Oncogene 41, 4211–4230 (2022).

33. Guelen, L. et al. Domain organization of human chromosomes revealed by mapping of nuclear lamina interactions. Nature 453, 948–951 (2008).

34. Robson, M. I. et al. Constrained release of lamina-associated enhancers and genes from the nuclear envelope during T-cell activation facilitates their association in chromosome compartments. Genome Res. 27, 1126–1138 (2017).

35. Mattick, J. S. et al. Long non-coding RNAs: definitions, functions, challenges and recommendations. Nat Rev Mol Cell Biol 24, 430–447 (2023).

36. Zeng, M. et al. DeepLncLoc: a deep learning framework for long non-coding RNA subcellular localization prediction based on subsequence embedding. Briefings in Bioinformatics 23, bbab360 (2022).

37. Li, J. et al. LncTar: a tool for predicting the RNA targets of long noncoding RNAs. Briefings in Bioinformatics 16, 806–812 (2015).

38. Crabbe, L., Cesare, A. J., Kasuboski, J. M., Fitzpatrick, J. A. J. & Karlseder, J. Human Telomeres Are Tethered to the Nuclear Envelope during Postmitotic Nuclear Assembly. Cell Reports 2, 1521–1529 (2012).

39. Lombardi, M. L. et al. The Interaction between Nesprins and Sun Proteins at the Nuclear Envelope Is Critical for Force Transmission between the Nucleus and Cytoskeleton*. Journal of Biological Chemistry 286, 26743–26753 (2011).

40. Gruenbaum, Y., Margalit, A., Goldman, R. D., Shumaker, D. K. & Wilson, K. L. The nuclear lamina comes of age. Nat Rev Mol Cell Biol 6, 21–31 (2005).

41. Stephens, A. D., Banigan, E. J., Adam, S. A., Goldman, R. D. & Marko, J. F. Chromatin and lamin A determine two different mechanical response regimes of the cell nucleus. MBoC 28, 1984–1996 (2017).

42. van Steensel, B. & Belmont, A. S. Lamina-Associated Domains: Links with Chromosome Architecture, Heterochromatin, and Gene Repression. Cell 169, 780– 791 (2017).

43. Dechat, T., Adam, S. A., Taimen, P., Shimi, T. & Goldman, R. D. Nuclear Lamins. Cold Spring Harb Perspect Biol 2, a000547 (2010).

44. Johnson, B. R. et al. A-type lamins regulate retinoblastoma protein function by promoting subnuclear localization and preventing proteasomal degradation. Proceedings of the National Academy of Sciences 101, 9677–9682 (2004).

45. Ivorra, C. et al. A mechanism of AP-1 suppression through interaction of c-Fos with lamin A/C. Genes Dev. 20, 307–320 (2006).

46. Neumann, S. et al. Nesprin-2 Interacts with α-Catenin and Regulates Wnt Signaling at the Nuclear Envelope*. Journal of Biological Chemistry 285, 34932–34938 (2010).

47. Spann, T. P., Goldman, A. E., Wang, C., Huang, S. & Goldman, R. D. Alteration of nuclear lamin organization inhibits RNA polymerase II–dependent transcription. Journal of Cell Biology 156, 603–608 (2002).

48. Wang, Y. et al. Lamin A/C-dependent chromatin architecture safeguards naïve pluripotency to prevent aberrant cardiovascular cell fate and function. Nat Commun 13, 6663 (2022).

49. Schreiber, K. H. & Kennedy, B. K. When Lamins Go Bad: Nuclear Structure and Disease. Cell 152, 1365–1375 (2013).

50. Chow, K.-H., Factor, R. E. & Ullman, K. S. The nuclear envelope environment and its cancer connections. Nat Rev Cancer 12, 196–209 (2012).

51. Swift, J. et al. Nuclear Lamin-A Scales with Tissue Stiffness and Enhances Matrix-Directed Differentiation. Science 341, 1240104 (2013).

52. Dechat, T. et al. Nuclear lamins: major factors in the structural organization and function of the nucleus and chromatin. Genes Dev. 22, 832–853 (2008).

53. Ikegami, K., Secchia, S., Almakki, O., Lieb, J. D. & Moskowitz, I. P. Phosphorylated Lamin A/C in the Nuclear Interior Binds Active Enhancers Associated with Abnormal Transcription in Progeria. Developmental Cell 52, 699–713.e11 (2020).

54. Ikegami, K., Secchia, S., Almakki, O., Lieb, J. D. & Moskowitz, I. P. Phosphorylated Lamin A/C in the Nuclear Interior Binds Active Enhancers Associated with Abnormal Transcription in Progeria. Developmental Cell 52, 699–713.e11 (2020).

55. Favorov, A. et al. Exploring Massive, Genome Scale Datasets with the GenometriCorr Package. PLoS Comput Biol 8, e1002529 (2012).

