## Supplementary material for "Distal Gene Expression Governed by Lamins and Nesprins via Chromatin Conformation Change": Supplemenrary file

### Supplementary Figures and figure legends:

### Supplementary Figure S1

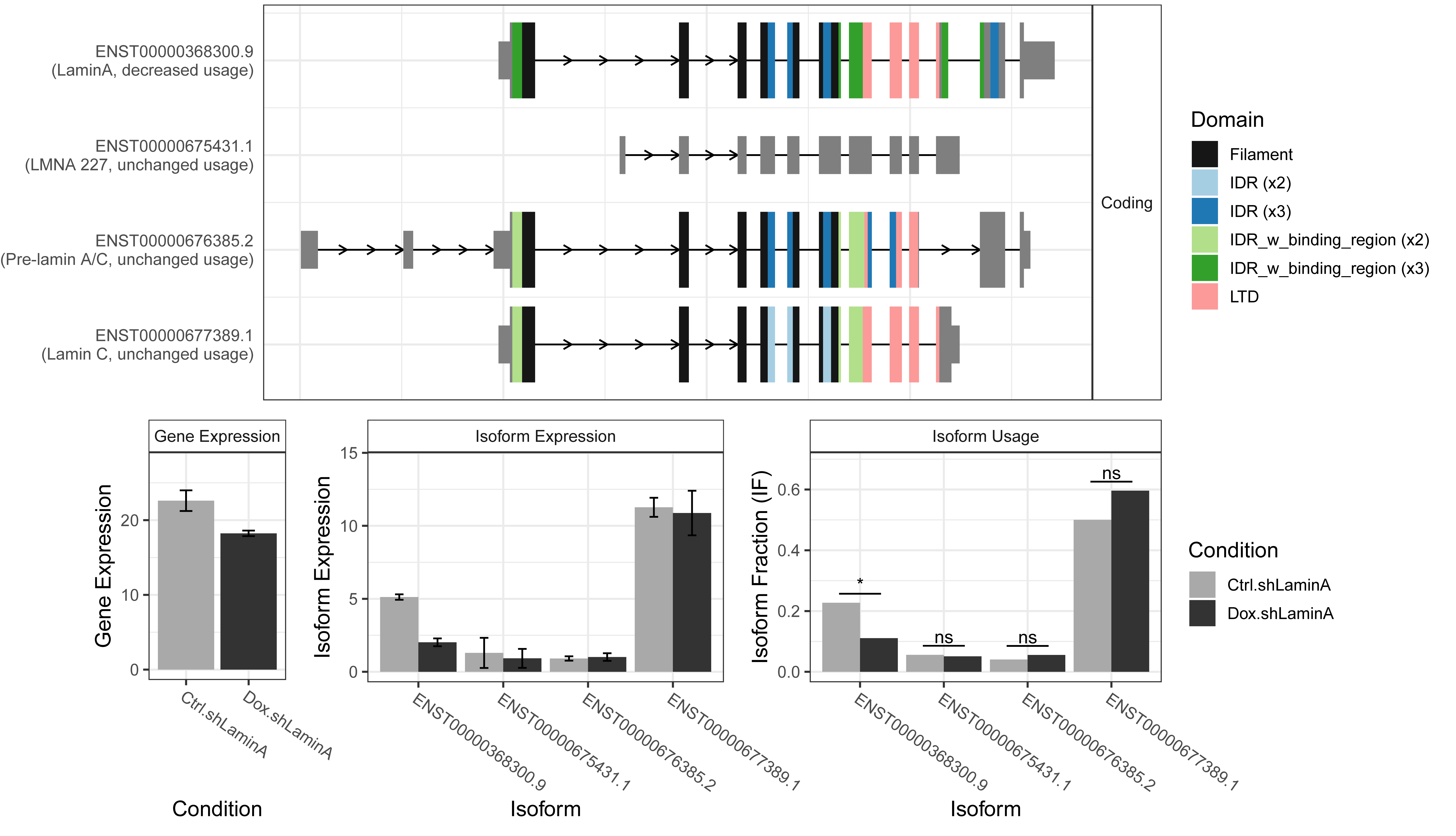

**Supplementary Figure S1.** Quantitative isoform switches of LMNA genes in control groups (Ctrl.shLaminA) versus doxycycline-treated groups (Dox.shLaminA). Using the IsoformSwitchAnalyzeR package, four isoforms of the LMNA gene were identified: lamin A (ENST00000368300.9), LMNA 227 (ENST00000675431.1), pre-lamin A/C (ENST00000676385.2), and lamin C (ENST00000677389.1). Our data demonstrated a significant reduction in lamin A isoform usage, while other isoforms remained unchanged. Statistical comparisons were performed using Fisher's exact test, and *p*-values were adjusted for multiple comparisons using the Benjamini–Hochberg procedure, with significance defined as *q*-value < 0.05.

**Supplementary Figure S2**

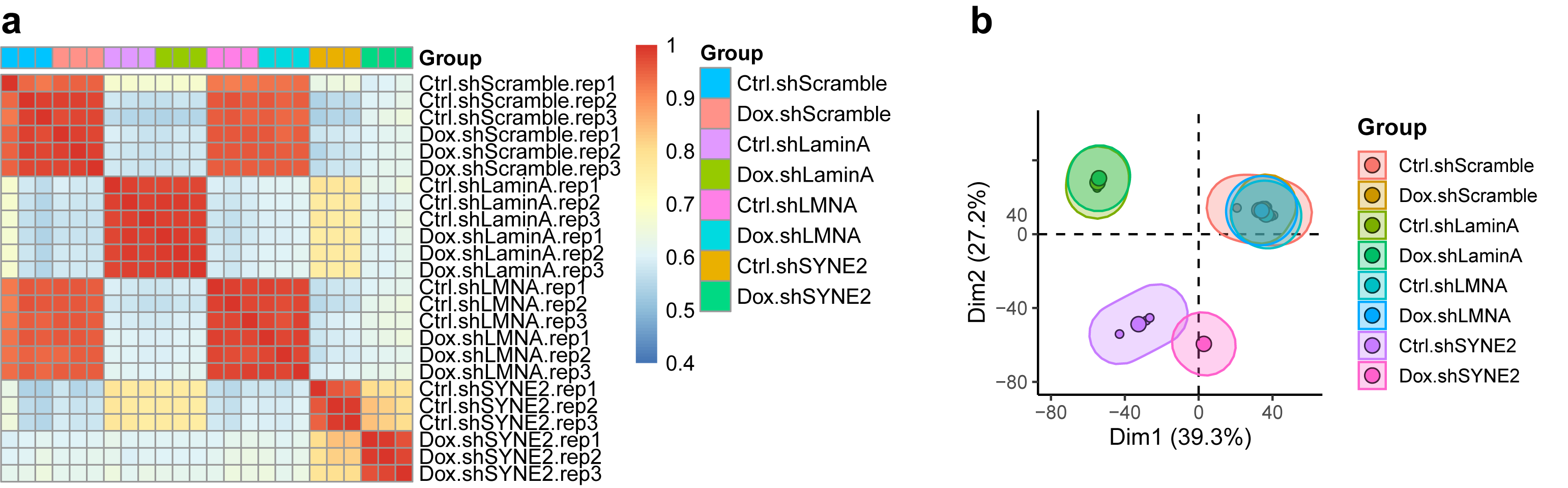

**Supplementary Figure S2. a)** Correlation heatmap depicting the correlation between samples, with each cell representing the relationship between a pair of samples. Red represents a strong correlation, while blue indicates a weaker correlation. **b)** Principal component analysis (PCA) plot of all samples, with the first principal component (Dim1) represented on the x-axis and the second principal component (Dim2) on the y-axis.

**Supplementary Figure S3**

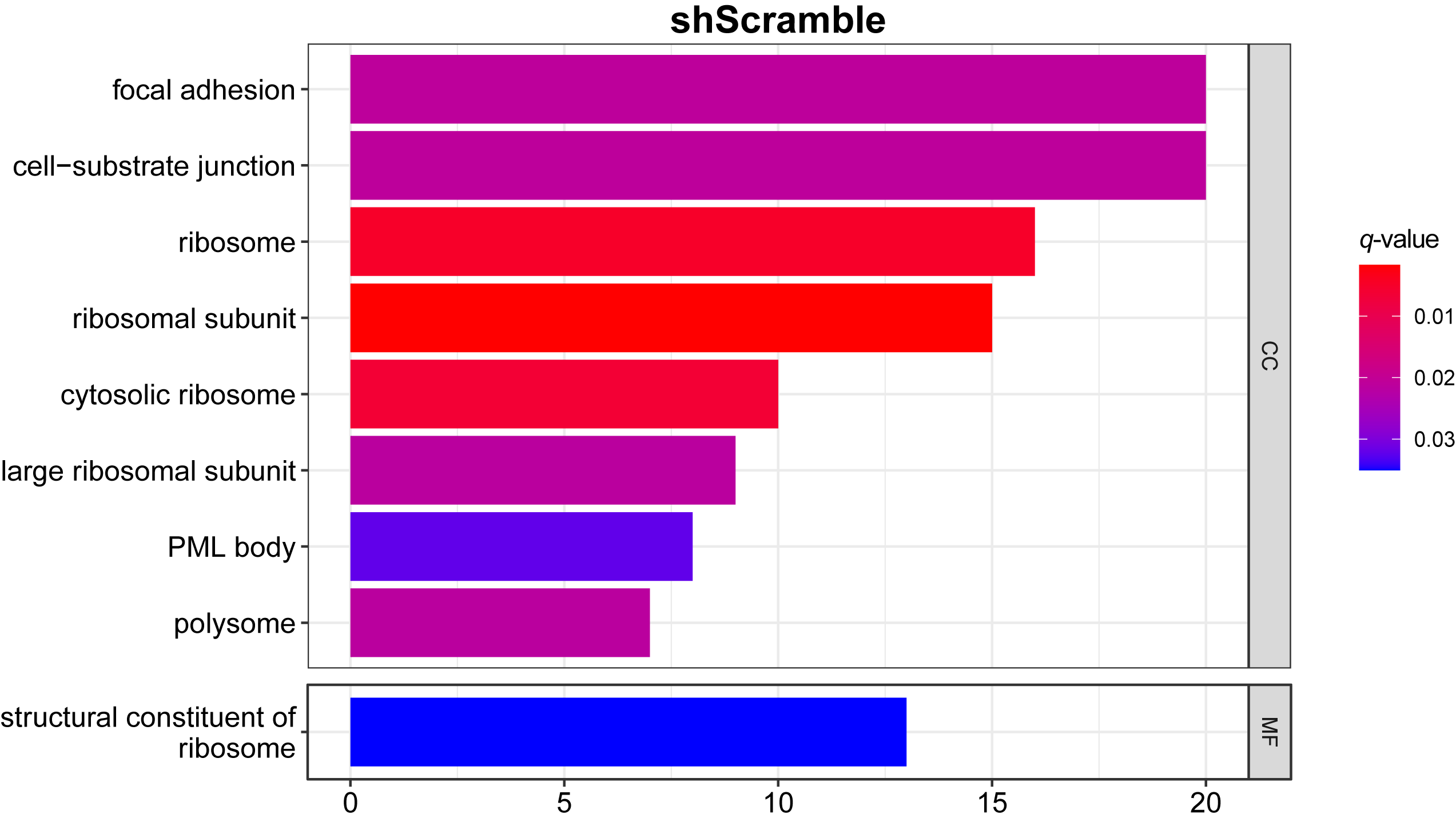

**Supplementary Figure S3.** Gene Ontology (GO) enrichment analysis of the differentially expressed genes (DEG) was performed for shScramble comparison. No significant biological process (BP) pathway was enriched.

**Supplementary Figure S4**

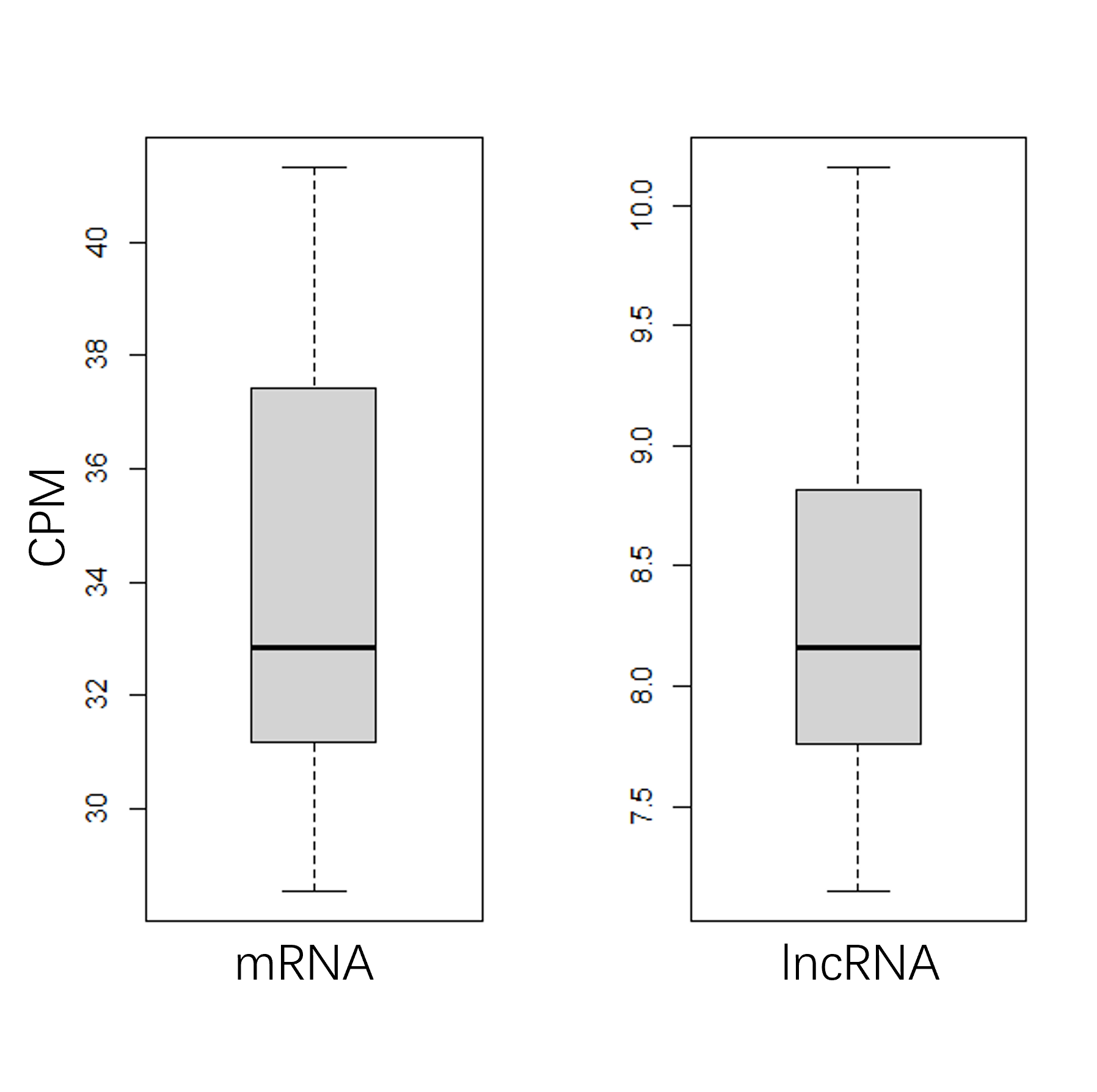

**Supplementary Figure S4.** Relative low expression of DE lncRNAs compared to DE mRNAs.
